# Astrocytic C-X-C motif chemokine ligand-1 mediates β-amyloid-induced synaptotoxicity

**DOI:** 10.1101/2021.09.22.458716

**Authors:** Beatriz Gomez Perez-Nievas, Louisa Johnson, Paula Beltran-Lobo, Martina M. Hughes, Luciana Gammallieri, Francesca Tarsitano, Monika A Myszczynska, Irina Vazquez-Villasenor, Maria Jimenez-Sanchez, Claire Troakes, Stephen B. Wharton, Laura Ferraiuolo, Wendy Noble

**Author notes:** **Corresponding authors:** Dr Beatriz Gomez Perez Nievas and Dr Wendy Noble, King’s College London, Department of Basic and Clinical Neuroscience, Maurice, Wohl Clinical Neuroscience Institute, 5 Cutcombe Road, LONDON. SE5 8RX. UK., or.

## Abstract

**Background:** Pathological interactions between β-amyloid (Aβ) and tau drive the synapse loss that underlies neural circuit disruption and cognitive decline in Alzheimer’s disease (AD). Reactive astrocytes, displaying altered functions, are also a prominent feature of AD brain. This large and heterogeneous population of cells are increasingly recognised as contributing to early phases of disease. However, the contribution of astrocytes to detrimental Aβ-tau interactions in AD is not well understood.

**Methods:** Mouse and human astrocyte cultures were stimulated with concentrations and species of human Aβ that mimic those in human AD brain. Astrocyte conditioned medium was collected and immunodepleted of Aβ before being added to rodent or human neuron cultures. Cytokines, identified in unbiased screens were also applied to neurons, including following the pre-treatment of neurons with chemokine receptor antagonists. Tau mislocalisation, synaptic markers and dendritic spine numbers were measured in cultured neurons and organotypic brain slice cultures.

**Results:** Conditioned medium from astrocytes stimulated with Aβ induces tau mislocalisation and exaggerated synaptotoxicity that is recapitulated following spiking of neuron culture medium with recombinant C-X-C motif chemokine ligand-1 (CXCL1), a chemokine we show to be upregulated in Alzheimer’s disease brain. Antagonism of neuronal C-X-C motif chemokine receptor 2 (CXCR2) prevented tau mislocalisation and synaptotoxicity in response to CXCL1 and Aβ-stimulated astrocyte secretions.

**Conclusions:** Our data indicate that astrocytes exacerbate tau mislocalisation and the synaptotoxic effects of Aβ via interactions of astrocytic CXCL1 and neuronal CXCR2 receptors, highlighting this chemokine-receptor pair as a novel target for therapeutic intervention in AD.

## Introduction

Synapse loss in neocortex and limbic areas is an early pathological feature of Alzheimer’s disease (AD) that correlates strongly with cognitive decline [1, 2]. The extent of synapse loss in AD brain cannot be fully accounted for by loss of neurons alone, implying that surviving neurons also lose synapses [3, 4]. Loss of connectivity between surviving neurons in AD brain damages the efficiency of neural systems, explaining the association between synapse loss and cognitive decline.

Bioactive soluble dimers, oligomers and to a lesser extent N-terminally extended Aβ peptides produced by cultured cells, transgenic rodent models of AD, or extracted from human AD brain, reduce dendritic spine number and disrupt synaptic function, long-term potentiation (LTP) and rodent cognition [5–9]. Yet, it remains to be established whether clearance of Aβ alone is sufficient to prevent synaptic degeneration in AD [10]. Missorting of modified forms of tau to both the pre-synapse and dendritic spines is also linked with synaptotoxicity in AD [11–14]. For example, phosphorylated tau allows Aβ-induced excitotoxicity at post-synapses by mediating phosphorylation of NR2B by fyn tyrosine kinase and disruptions of NR2B-PSD-95 interactions [11], while at pre-synapses, phosphorylated tau binds synaptic vesicle proteins including synaptogyrin-3 to anchor vesicles and impair their release during sustained neuronal activity [13, 14]. We previously reported an association of synaptic tau with dementia in AD. Phosphorylated and oligomeric forms of tau were redistributed from the cytosolic compartment into synaptoneurosomes in cases with typical AD pathology, synapse loss and dementia, but not in so-called “mismatches” who showed a similar burden of AD pathology but preserved synaptic protein levels and cognition [15]. In addition, we saw increased astrocyte reactivity, as indicated with GFAP immunolabelling, in those with AD and dementia relative to mismatch cases [15].

Astrocytes are an intrinsic component of synapses [16] and affect synaptic activity by regulating the availability of glutamate, GABA, ATP, and glucose [17–20]. As such, healthy functioning astrocytes are critical for synaptic transmission [21], neural circuit maintenance [22] and long-term potentiation [23].

In AD, and particularly in association with elevated Aβ or amyloid plaques [24–26], astrocytes become “reactive” leading to changes in their morphology, molecular fingerprint and function [27, 28], and reactive astrocytes are often closely associated with increased disease severity and cognitive decline [29].

However, the specific contribution of reactive astrocytes to AD pathogenesis remains unclear, with some suggesting that reactive astrocytes lose synaptic and neuronal support functions [30] or that astrocytes undergo cellular senescence in AD [31]. Others report neurotoxic effects of reactive astrogliosis, including those mediated by inflammatory astrocytic secretions [32–34]. Particularly in response to repeated insult, systemic or secondary inflammation, astrocytes show exaggerated production and secretion of pro-inflammatory cytokines including interleukin (IL)-1beta, IL-6 and chemokines such as CXCL1 [35, 36].

Reactive astrocytes may also be neuroprotective, with recent reports showing that astrocytic IL-3 signals to microglia to promote their phagocytosis of aggregated tau and Aβ [37]. This divergence in astrocytic response may be at least partly related to astrocytic heterogeneity in different brain regions and in response to different types of acute injury [30, 38, 39]. Here, we sought to better understand the contribution of astrocytes to synaptotoxic Aβ-tau interactions in AD by exposing rodent and human astrocytes to concentrations and species of human Aβ that replicate those found in human AD brain.

## Methods and Materials

All animal work was conducted in accordance with the UK Animals (Scientific Procedures) Act 1986 and the European Directive 2010/63/EU under UK Home Office Personal and Project Licenses and with agreement from the King’s College London (Denmark Hill) Animal Welfare and Ethical Review Board.

### Rodent primary neural cell cultures

Primary astrocytic cultures were prepared from the cortex of wild type CD1 mice on postnatal day 1-3 as previously described [40]. The astrocytes were seeded onto a poly-D-lysine (PDL 10 μg/mL) precoated T75 flasks and maintained in culture for 7– 10 days in a humidified CO_2_ incubator at 37 °C with shaking at 200 rpm overnight on days 3 and 7 to remove remaining microglia and oligodendrocytes. Astrocyte-enriched cultures were trypsinized using TrypLE (ThermoFisher Scientific) and replated on PDL precoated 6- or 12- wells plates. 24 hours before treatment, growth medium (high glucose DMEM with glutamax, 10% fetal bovine serum, 100 units/ml penicillin and 100 μg/ml streptomycin) was changed to Neurobasal-B27 serum-free medium.

Primary neurons were obtained from cerebral cortex of CD1 embryos at embryonic day 15, as previously described [41]. Neurons were plated at a density of 6×10^5^ viable cells/35mm^2^ on glass-bottomed dishes (MatTek Corporation, Ashland, MA, USA), 96-well plates (BD Falcon) or 12-well Nunc™ plates (ThermoFisher Scientific) previously coated with PDL (10 μg/ml) for at least 1 h at 37°C. Cultures were maintained at 37°C in 5% CO_2_, in neurobasal medium with 2% B27 nutrient, Glutamax, penicillin (100 units/ml) and streptomycin (100 μg/ml). At 13 days *in vitro* (DIV) neurons, transfected at 5-7 DIV with the plasmid peGFP-N1 (Clontech, Mountain View, CA) using lipofectamine 2000 (ThermoFisher Scientific), were treated for 24 hours and at 14 DIV dendritic spines were imaged or cells were fixed or lysed.

To obtain Aβ-containing medium (transgenic conditioned media or TGCM), neurons from Tg2576 mice overexpressing human APP containing the Swedish mutation K670NM671L were cultured, and after 14 DIV medium from healthy neurons was collected [42]. Media collected from 14 DIV wild-type neurons cultured from littermates (wild type conditioned media or WTCM) was used as a control. The genotype of the animals was determined by polymerase chain reaction on DNA obtained from the embryos.

For co-culture experiments, primary astrocytes were plated on cell-culture inserts (0.4 μm pore membrane, Falcon, Corning, Corning, NY, USA) that allow the passage of small molecules between cells and culture medium, and neurons were plated on 6 well plates, as described above. Astrocytes were treated with TGCM or WTCM for 24 hours, the medium was removed (TGCM astro and WTCM astro), the cells washed with PBS and inserts placed on top of cultured neurons for another 24 hours. There was no direct contact between neurons and astrocytes.

### Human astrocyte and neuronal cell cultures

All skin biopsy donors provided informed consent before sample collection (University of Sheffield, Study number STH16573, Research Committee reference 12/YH/0330). Skin fibroblasts from three control subjects were used (Table 2). These were reprogrammed as previously described [43]. Once induced, neuronal progenitor cells (iNPC) cultures were established, and the medium was switched to NPC proliferation media consisting of DMEM/F12 (1:1) GlutaMax, 1% N2, 1% B27, and 40 ng/ml FGF2. iNPC-derived astrocytes were produced as previously described [43–45]. Briefly, iNPCs were switched to astrocyte proliferation media, DMEM (Thermo Fisher Scientific), 10% FBS (Life science production, Bedford, UK), 0.2% N2 (Thermo Fisher Scientific). Cells were differentiated in 6-well plates coated with human fibronectin for 7 days and were used at a matched number of passages.

Lund human mesencephalic (LUHMES) neuronal precursors were grown in T75 flasks in proliferation medium (DMEM/F12 GlutaMAX™ supplement medium (Thermo Fisher Scientific), N2 supplement (Thermo Fisher Scientific) and 40 ng/ml recombinant basic fibroblast growth factor (FGF) (Peprotech). To allow high content imaging on the Opera Phenix microscope, non-differentiated LUHMES were transduced with GFP-expressing lentiviral particles (LV-GFP), with GFP expression under the control of a PGK promoter as previously described [46]. When cells were 50–60% confluent they were differentiated by adding differentiation media consisting of DMEM/F12 GlutaMAX™ supplement medium, N2 supplement and 1 μg/ml tetracycline. After two days, LUHMES were trypsinized and replated onto 96 wells plates for a further 3 days before experimentation.

### Postmortem human brain

Post-mortem human prefrontal cortex (Brodmann area 9; BA9) from control and clinically and pathologically confirmed cases of Alzheimer’s disease were obtained from the Neurodegenerative Diseases Brain Bank, King’s College London. All tissue collection and processing were carried out under the regulations and licensing of the Human Tissue Authority, and in accordance with the UK Human Tissue Act, 2004. Samples were collected from control cases (Braak stage 0, n = 4) and those with mild AD neuropathology (Braak stages I-II, n= 7), moderate AD neuropathology (Braak stages III-IV, n= 10), and severe AD neuropathology (Braak stages V-VI, n= 6) (Table 1). There were no significant differences in age, gender or post-mortem delay between groups.

**Table 1:**
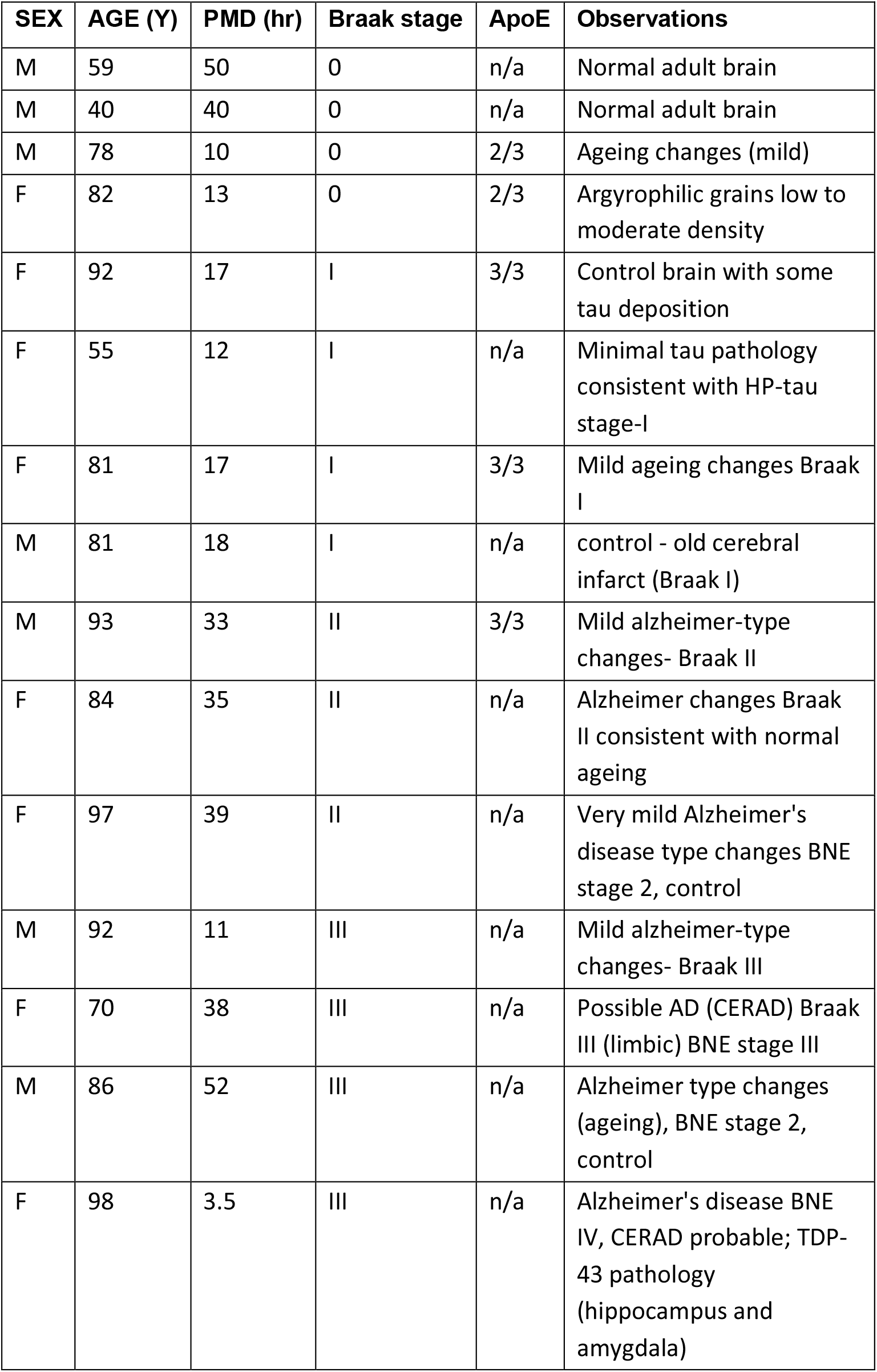
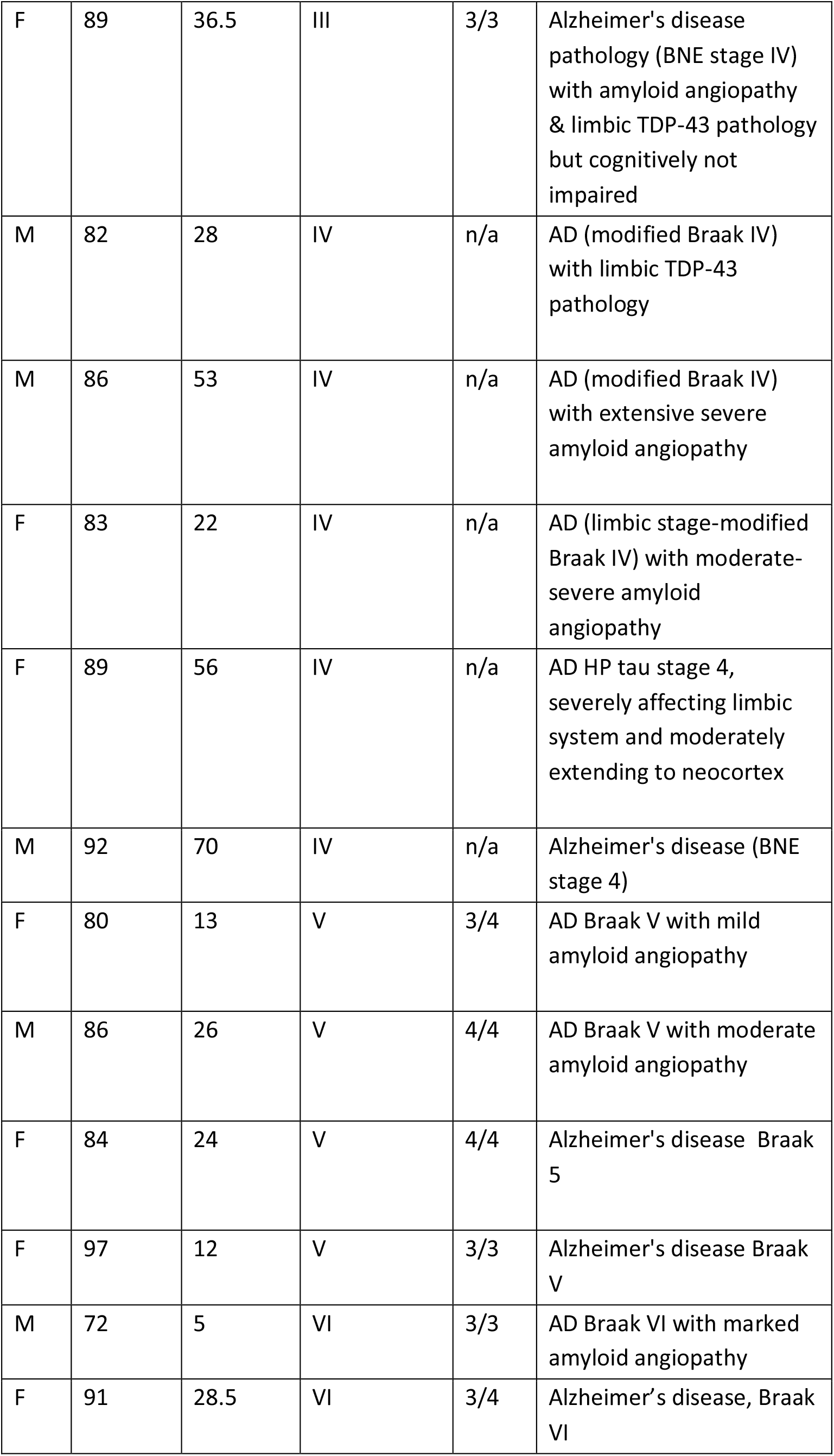
Human postmortem samples from BA9 prefrontal cortex, that were used in this study. Age is shown in years (y), postmortem delay (PMD) in hours (h), sex and Braak stage. Braak stage 0 cases had some phospho-tau but did not meet Braak stage I criteria and were used as controls.

**Table 2:**
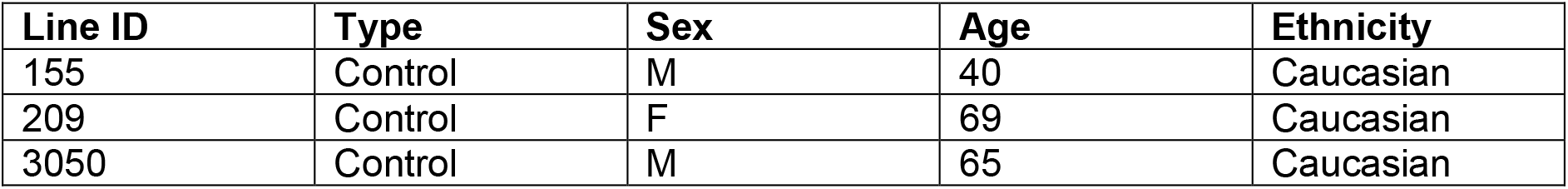
Characteristics of human donors from whom fibroblasts were obtained. Line ID, type (disease or control), sex, age in years and ethnicity are shown.

### Mouse organotypic brain slice culture

Organotypic brain slice cultures were prepared from P9 wild-type CD1 mice and cultured as described in Croft et al. [47]. Slices were treated after two weeks in culture with WTCM or TGCM as described for astrocytes.

### Cell and slice culture treatments

The concentration of Aβ in TGCM was determined by ELISA (ThermoFisher Scientific) following the manufacturer’s instructions and was adjusted to 2000 pM Aβ 40/200 pM Aβ 42 prior to use. An equivalent volume of WTCM was used as control. To remove Aβ from the medium of astrocytes challenged with TGCM, media was incubated with dynabeads protein G (ThermoFisher) bound with 6E10 antibody (COVANCE) for 1hour at 4C. Beads were separated from medium using a magnetic stand to remove Aβ-6E10 complexes. The efficacy of immunodepletion was assessed by measuring levels of Aβ before and after 6E10 immunodepletion by ELISA and western blot (Supplementary Fig. 1).

To test the direct effects of the cytokine CXCL1 on synapse/neuronal health, DIV12-13 neurons were pre-treated for 20 minutes with 40 ng/ml of the CXCR2 (CXCL1 receptor in neurons) antagonist SB225002 (Tocris Bioscience), prior to the addition of 40 nM recombinant mouse CXCL1 (R&D Systems) or vehicle for a further 24 hours.

### Isolation of synaptoneurosomes

Total, cytosolic and synaptic fractions were isolated from human brain tissues and mouse organotypic brain slice cultures as previously described [15, 48]. Protein concentrations were determined using a bicinchoninic acid (BCA) protein concentration assay kit (Pierce) according to the manufacturer’s instructions and were adjusted to the same concentration for all sample*s* by adding homogenisation buffer. Tau content in total, cytosolic and synaptoneurosomal fractions was examined for organotypic brain slice culture experiments. The cytosolic fraction of postmortem human brain was used for cytokine arrays.

### Measurement of dendritic spine density, neuron complexity and tau mislocalisation

NeuronStudio software (CNIC, Mount Sinai School of Medicine) was used for dendritic spine analysis. Spine density, or number of neuritic protrusions was defined as number per micrometer of dendrite/neurite length. Dendritic spine densities were calculated from approximately 20 neurons/condition for each biological replicate. Neuronal morphology of 14 DIV neurons transfected with the plasmid peGFP-N1 (Clontech, Mountain View, CA) was examined using high-resolution confocal digital images obtained from an Opera-Phenix microscope (Perkin-Elmer). Neuronal complexity was analysed using Harmony software which gives measures of total and maximum neurite length, number of roots, nodes, segments and extremities. The number of neurites per cell containing missorted tau was assessed with Harmony^TM^ software using an algorithm that identifies neurite segments labelled with MAP2, that contain tau.

The amount of lactate dehydrogenase (LDH) in the media of cultured neurons was determined as a measure of neuron health, using an LDH Cytotoxicity Kit from Thermo Fisher Scientific, according to the manufacturer’s instructions.

### Cytokine arrays

The cytoplasmic fraction of human postmortem brain homogenates and astrocyte culture media were used to determine cytokine content using Mouse or Human Proteome Profiler arrays (Mouse Cytokine Array Panel A and Human Cytokine Array, R&D Systems), according to the manufacturers’ instructions. Positive and negative control spots included were used to allow quantitative analysis of cytokine levels. Results were expressed as percentage change compared to controls.

### SDS-PAGE and western blotting

Cells were washed with PBS and directly lysed into PBS containing sample buffer (NuPAGE LDS Sample Buffer 4X, Invitrogen), reducing agent (NuPAGE Sample Reducing Agent 10X, Invitrogen), protease inhibitor (complete Mini EDTA-Free Protease Inhibitor Cocktail, Roche, Basel, CH) and phosphatase inhibitor (PhosSTOP, Roche) cocktails.

Equal amounts of protein were separated on 10% or 4–12% SDS-PAGE gels (Thermo Fisher Scientific) by electrophoresis, transferred to 0.45 μm nitrocellulose membranes (Millipore, MA, USA), and immunoblotted as described previously (Glennon et al., 2020). The primary antibodies used were against PSD-95 (Rabbit IgG, #3409, Cell Signalling 1/500), synapsin-1 (Rabbit IgG, #BV-6008, Enzo 1/500), caspase-3 (Rabbit IgG, #9661, Cell Signalling 1/500), GFAP (Mouse IgG, Z0334, DAKO Ltd, 1/500), phosphoNFkB (Rabbit IgG, #3031, Cell Signalling, 1/1000), β-amyloid-6E10 (Mouse IgG, SIG-39320, 1/200-1/500), lipocalin-2/NGAL (Mouse IgG, #AF1857, 1/500), tau (Rabbit IgG, #A0024, DAKO Ltd, 1/5000), PHF1 (Mouse IgG, P. Davies, 1/2000), GAPDH (Mouse IgG, #32233, SantaCruz Biotechnology, 1/500), β-actin (Mouse IgG, #ab8226, 1/1000), α-tubulin (Rabbit IgG, #ab18251, 1/1000). Bound antibodies on membranes were imaged using an Odyssey CLX instrument (LI-COR Biosciences) and analysed using Image Studio software LI-COR Biosciences.

### Immunocytochemistry

Cells were fixed for 10 minutes in 4% formaldehyde and 4% sucrose in PBS and then 10 minutes in pre-chilled methanol. Cells were simultaneously permeabilized and blocked using PBS containing 2% normal goat serum and 0.1% Triton-X-100 for 1 hour at room temperature and immunolabelled according to previously published protocols [49]. Primary antibodies were against tau (Rabbit IgG, #A0024, DAKO, 1/500), MAP2 (Mouse IgG, #GTX8266, GeneTex, 1/200), and GFAP (Rabbit IgG, #Z0334, DAKO, 1/100). 96 wells plates were directly imaged on the Perkin Elmer’s Opera Phenix microscope and glass-bottomed dishes on the Nikon Ti-E 3 camera inverted microscope after adding a drop of Dako fluorescent mounting media to each well.

### Statistical analysis

Data were analysed using GraphPad Prism. After performing a Shapiro-Wilk normality test, most data were analysed using one-way ANOVA followed by Tukey *post hoc* test or Student’s t-test (GraphPad Prism 7 Software, Graphpad Software, La Jolla, CA, USA). Results were considered statistically significant when P<0.05. For neurons treated with astrocyte conditioned media and SB225, a two-way ANOVA followed by post hoc tests was performed (see figure legends). Data are shown as mean ± standard error of the mean (SEM).

## Results

### Conditioned medium from Aβ stimulated astrocytes is synaptotoxic

Primary neurons cultured from Tg2576 mice release human Aβ into the culture medium in an approximately 1:10 ratio of Aβ42:Aβ40 as in human AD brain (Fig. 1A). Exposure of naïve neurons to physiological concentrations of Aβ (200pM Aβ42, 2000pM Aβ40) in medium from Tg2576 mice (TGCM) induces a decrease in the number of dendritic spines compared to neurons treated with medium from wild type littermates (WTCM) [41, 50, 51]. This finding was reproduced here (Fig. 1B,C) with spine numbers in TGCM-stimulated neurons being reduced by approximately 36% to 66.42+/− 6.233 % relative to WTCM conditions (p<0.001). Notably, spine loss was exacerbated (reduction of 51.60+/− 5.26 SEM% relative to WTCM) when neurons were exposed to medium from astrocytes stimulated with TGCM (TGCM astro) compared to those treated with WTCM (WTCM astro). Importantly, this effect is still observed after immunodepleting Aβ from the medium using 6E10 (TGCM astro-ID) (Fig. 1B,C; Supplementary Fig. 1), demonstrating that soluble factors released by astrocytes mediate this heightened synaptotoxicity.

**Figure 1:**
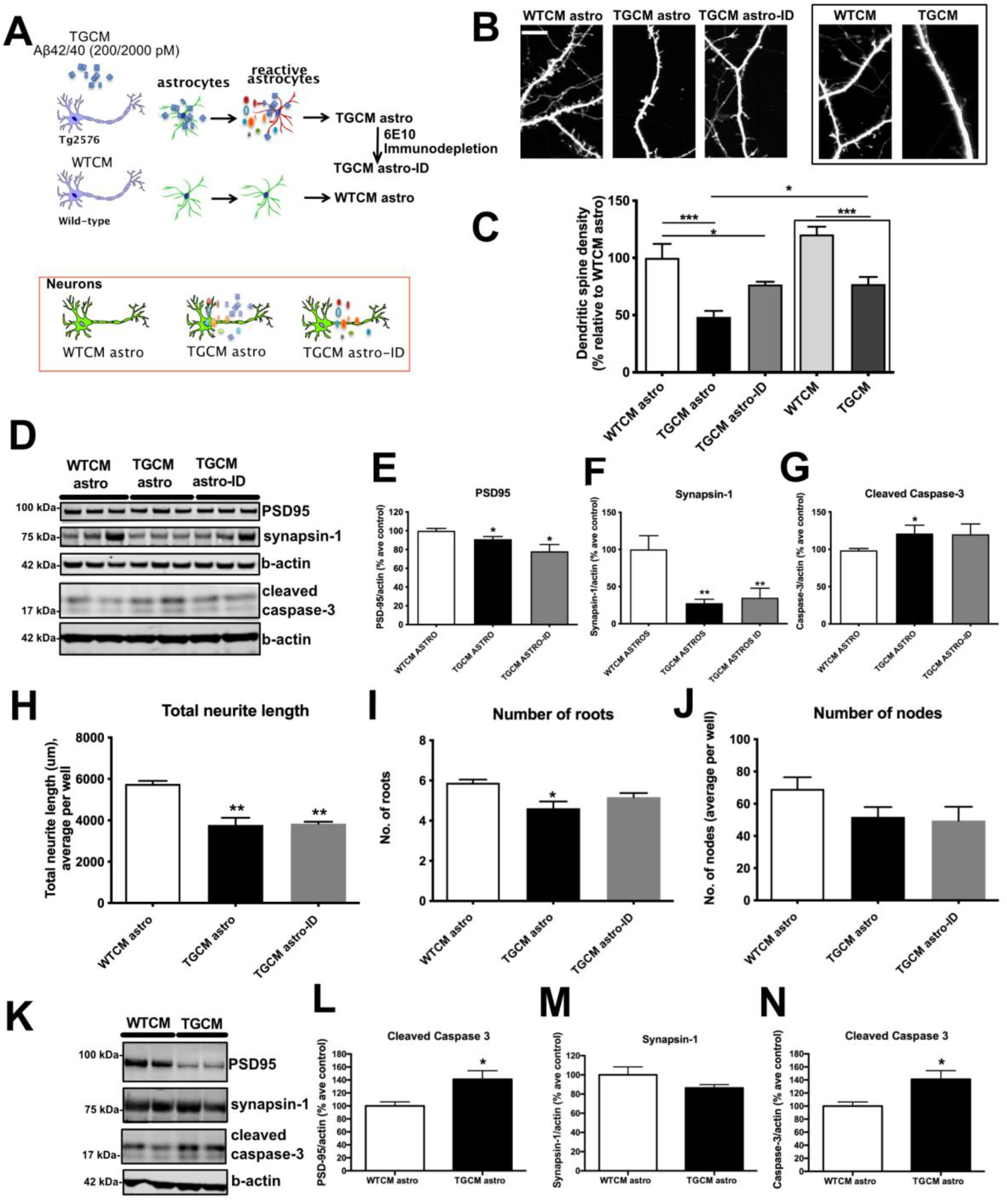
Conditioned medium from Aβ-stimulated astrocytes exaggerates the damaging effects of Aβ at synapses. A) Schematic describing the experimental approach. Conditioned medium was collected from primary cortical neurons from wild-type or Tg2576 mice (WTCM; TGCM) and applied to astrocytes. The conditioned medium from stimulated astrocytes was collected, (WTCM astro and TGCM astro) and in some experiments Aβ was immunodepleted from the medium of TGCM treated astrocytes (TGCM astro-ID). These media were added to naïve WT primary neurons. B) Dendritic spine density was measured in neurons transfected at 7DIV with GFP and treated with WTCM astro, TGCM astro and TGCM astro-ID or directly with WTCM and TGCM at 13DIV for 24 hours. Representative images are shown. Scale bar is 10μm. C) Quantification of spine density in 10-20 neurons per experimental replicate treated with conditioned medium. TGCM astro caused increase dendritic spine loss relative to treatment with TGCM directly, and this is maintained when Aβ is immunodepleted from the medium. n= 8. D) Neuronal lysates were immunoblotted using antibodies against PSD-95 (95 kDa), synapsin-1 (77kDa), cleaved caspase-3 (17-19kDa) and β-actin (42kDa). Representative blots are shown. Western blot band intensities were quantified and showed that TGCM astro induces decreases in the synaptic markers E) PSD-95, F) synapsin-1 and increases in G) cleaved (active) caspase-3, that is maintained when Aβ was immunodepleted (TGCM astro-ID). n= 8. Neuronal complexity as a measure of neuron health was analysed for all cells in three wells per experimental replicate using Harmony software. This showed H) reduced neurite length, and fewer I) number of roots and J) nodes following treatment with TGCM astro and TGCM astro-ID, indicating that astrocytic secretions are damaging to neurons (n= 5). Astrocytes were cultured and treated in cell culture inserts, the medium removed by washing and the astrocytes added to neuron cultures. K) Representative western blots of neuronal lysate immunoblots from co-culture experiments. Western blot data was quantified and showed reduced L) PSD-95 and M) synapsin-1 and N) increased cleaved caspase-3 when neurons were co-cultured with astrocytes that had previously been exposed to TGCM (n= 6). Data on graphs is mean +/− SEM and is shown relative to values for neurons exposed to astrocytes treated with WTCM. *p<0.05, **p<0.01.

A reduction in the levels of synaptic proteins including PSD95 and synapsin-1 was found in neurons stimulated with medium from TGCM-stimulated astrocytes compared to medium from WTCM exposed astrocytes (Fig. 1D-F). This was accompanied by increases in levels of cleaved (active) caspase-3 (Fig. 1D,G), an executioner caspase which also plays various non-apoptotic roles in neurons including modulation of synaptic functions [52]. The changes in synaptic proteins and cleaved caspase-3 are induced by astrocytic secretions since they are still observed after Aβ is immunodepleted from the astrocyte-conditioned TGCM (TGCM astro-ID; Fig. 1D-G). Similarly, neurons treated with medium from TGCM-stimulated astrocytes showed indicators of poor health including reduced neuronal complexity, significant reductions in total neurite length and number of roots, alongside apparent decreases in other parameters that measure neuritic ramification (Fig. 1H-J; Supplementary Fig. 2).

To validate this data in more physiological co-culture conditions, astrocytes were grown on cell culture inserts and stimulated with TGCM or WTCM. The culture medium was replaced to remove traces of Aβ and the stimulated astrocytes co-cultured with neurons (Fig. 1K). Under these conditions, astrocytic secretions were also found to disrupt synapses indicated by significantly reduced PSD-95 (p<0.05), apparent reductions in synapsin-1 levels, and increases in cleaved caspase-3 (p<0.05) (Fig. 1K-M) in the absence of elevated lactate dehydrogenase (LDH) release (not shown). This confirms that factors released by astrocytes in response to concentrations of human Aβ, similar to that found in AD brain, compromise synaptic and neuronal health without causing overt neurotoxicity.

Despite recent studies indicating significant conservation between human and mouse astrocytes, there are species-specific differences in their response to stressors [53]. Therefore, human iNPC-astrocytes, which retain age-related features [54], were stimulated with TGCM or WTCM, the astrocyte conditioned medium collected and immunodepleted of Aβ with 6E10, prior to its addition to human post-mitotic neurons that were differentiated from LUHMES cells. LUHMES are a fetal human mesencephalic cell line conditionally immortalised with a myc transgene which become post-mitotic mature neurons when transgene expression is suppressed [55]. Similar to findings with mouse cells, medium from TGCM-exposed human astrocytes (TGCM h-astro) induced a significant loss of neuritic protrusions (Fig. 2A-B, p<0.001) and reductions in features of neuronal health and complexity relative to WTCM h-astro (p<0.05, Fig. 2C-D; Supplementary Fig. 3), that were retained in the absence of Aβ (TGCM h-astro-ID, p<0.05 for all).

**Figure 2:**
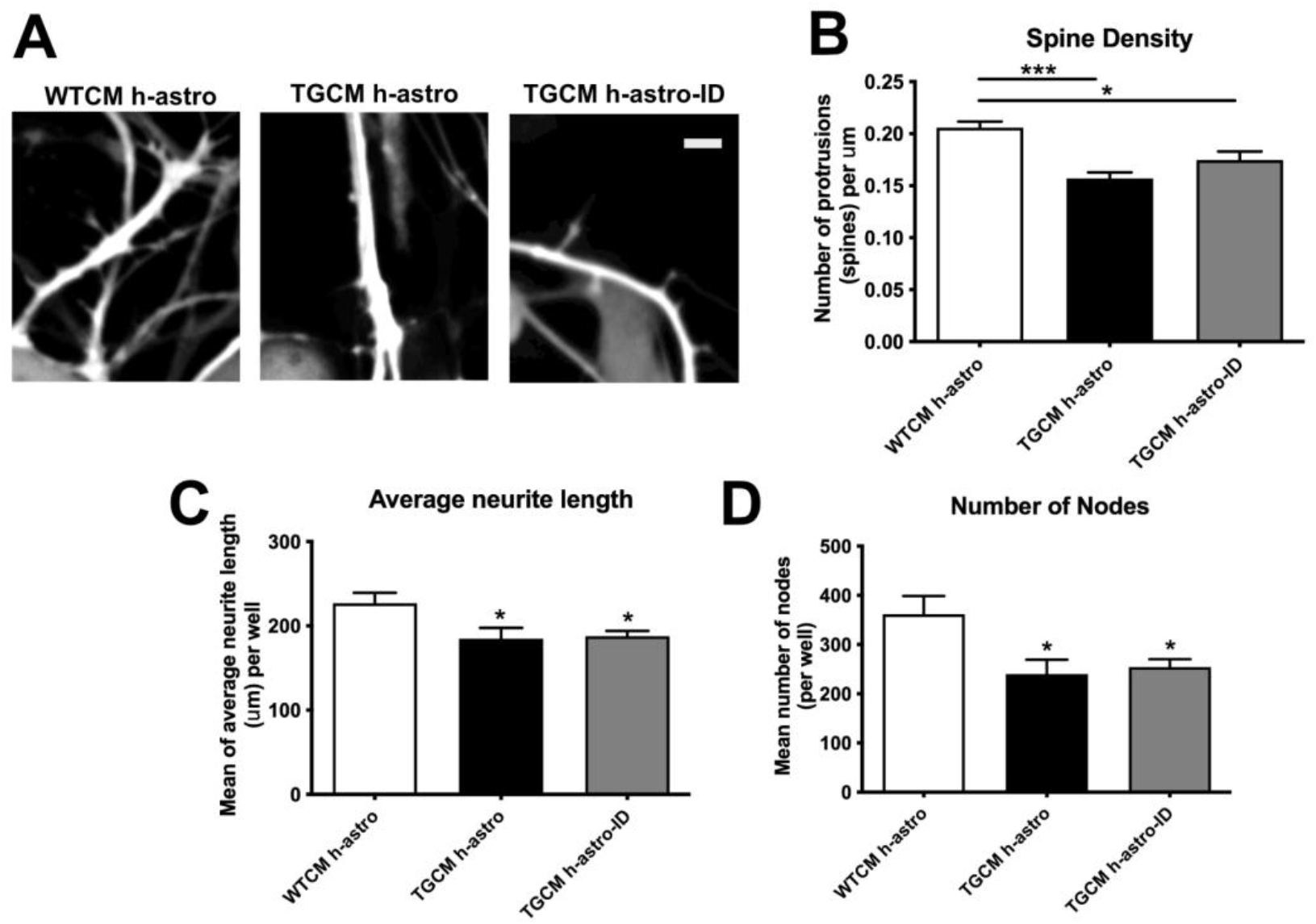
iNPC-astrocytes stimulated with human Aβ release synaptotoxic secretions. Human iNPC-astrocytes were exposed to WTCM and TGCM at 7 days post-differentiation for 24 hours. WTCM h-astro and TGCM h-astro was collected and in some cases Aβ was immunodepleted from astrocyte medium (TGCM h-astro-ID). A) The number of neuritic protrusions per μm in LUHMES transfected pre-differentiation GFP were imaged as proxy for dendritic spines in 20 cells per experimental repeat (indicated by arrow heads). Scale bar is 5μm. B) Quantification of protrusion number per μm shows reductions in the presence of TGCM h-astro relative to WTCM h-astro that is maintained when Aβ is removed from astrocyte conditioned medium by immunodepletion (TGCM h-astro-ID). Measures of neuron health across three wells per experimental repeat showed C) shorter average neurite length and D) fewer number of nodes following treatment with TGCM h-astro and TGCM h-astro-ID, indicating that human astrocytes secrete factors that are damaging to neurons (n= 3). Data on graphs is mean +/− SEM. *p<0.05, ***p<0.001.

### Astrocyte-mediated synaptotoxicity in response to Aβ is related to tau mislocalisation

Direct effects of Aβ on synapse and neuron health are tau-dependent [11, 54] and related to damaging effects of mislocalised tau at post-synapses [11, 12] and pre-synapses [13, 14]. We found that tau localisation was altered in response to medium from TGCM astro, with tau showing increased neuritic localisation (Fig. 3A-B). Similar findings were observed when organotypic brain slice cultures which contain all neural cell types, were treated with TGCM. The amount of tau phosphorylated at Ser396/404 (PHF1) was increased in the synaptic compartment when compared to slices treated with WTCM (Fig 3C-E).

**Figure 3:**
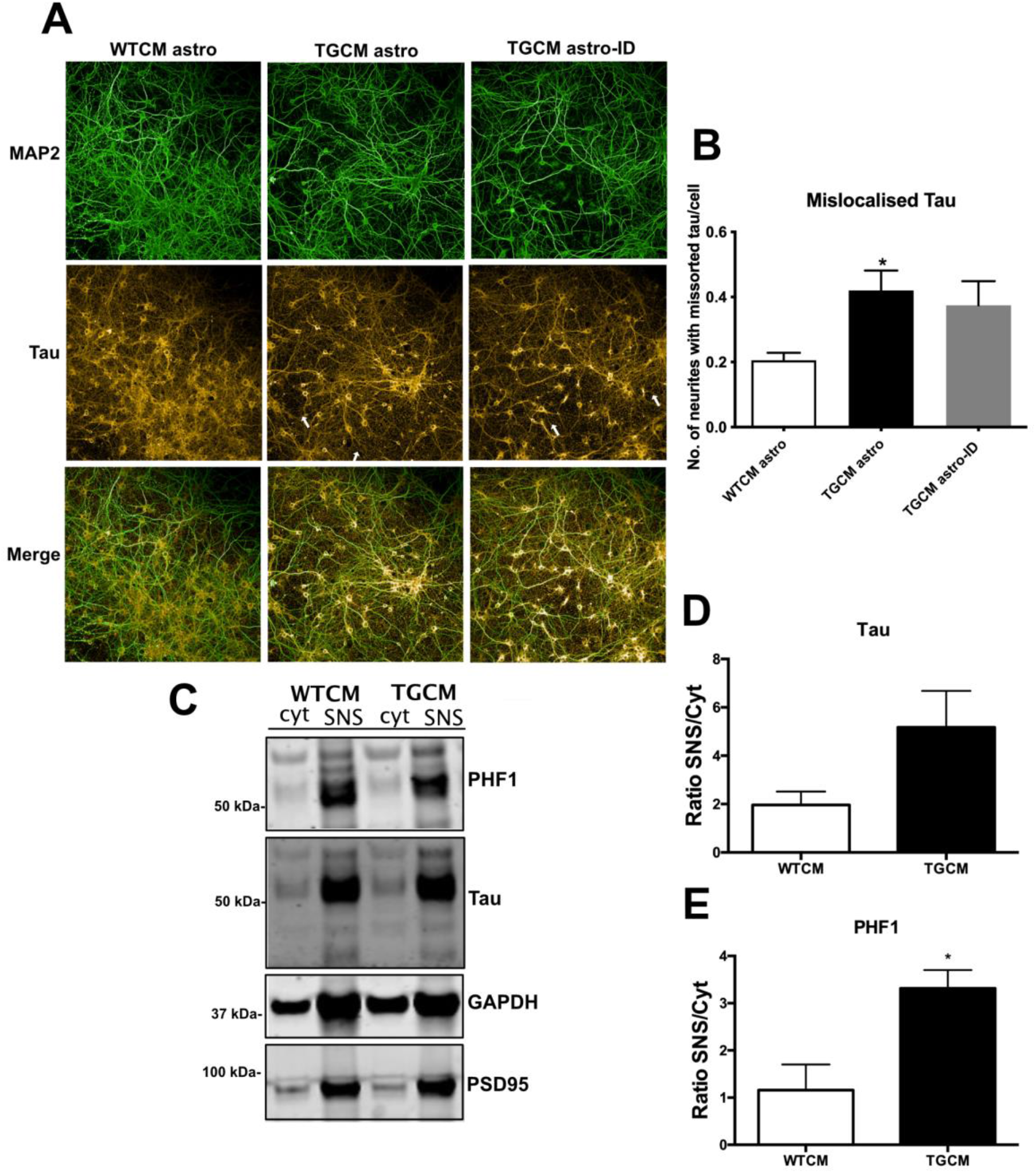
Astrocyte-mediated synaptotoxicity in response to Aβ is accompanied by tau mislocalisation. A) 14DIV neurons treated for 24 hours with WTCM astro, TGCM astro or TGCM astro-ID were fixed and immunolabelled with antibodies against MAP2 (green) and tau (far red). Exposure to TGCM astro and TGCM astro induced increased tau localisation in neurites with white arrows indicating beading of these neurites. Scale bar is 100μm. B) The number of MAP2 labelled neurites per cell containing missorted tau was quantified, showing increased tau mislocalisation upon exposure to conditioned medium from astrocytes exposed to Aβ (TGCM astro and TGCM astro-ID) (n= 4). C) To confirm that Aβ induces tau mislocalisation in the presence of other neural cell types, 14DIV organotypic brain slice cultures prepared from CD1 mice were treated with WTCM or TGCM for 24 hours. The cytosolic fraction (Cyt) and synaptoneurosomes (SNS) were extracted and immunoblotted with antibodies against PSD-95, total tau and tau phosphorylated at Ser396/404 (PHF1). GAPDH was used as a loading control. PSD-95 accumulates in the synaptic fraction showing successful SNS extraction. The abundance of D) tau and E) PHF1 in synaptoneurosomes relative to cytosolic was determined. Phosphorylated tau was mislocalised from the cytosol to synapses following exposure to Aβ (n= 3). Data on graphs is mean +/− SEM. *p<0.05.

### Astrocytic inflammatory phenotypes are induced by Aβ

Although microglia are considered the resident immune cell in the brain, the contribution of astrocytes to neuroinflammatory processes in AD is increasingly recognised [25, 35, 36]. We observed increased levels of GFAP, and other markers of astrocyte reactivity including lipocalin-2 (Lcn2), in astrocytes stimulated with TGCM relative to WTCM (Fig. 4A-G). Lcn2 was identified as a pan-reactive astrocyte marker in gene expression analyses [38] and we previously showed significant elevations in astrocytic lcn2 levels upon their exposure to oxysterol mixtures that mimic their composition in AD brain [34]. These changes reflect the induction of inflammatory signalling pathways in astrocytes since we observed increased phosphorylation of NFκB (p65) upon exposure of astrocytes to Aβ-containing medium relative to control (WTCM) conditions (Fig. 4C, F). We examined cytokine release from astrocytes using cytokine arrays which allowed unbiased measurement of a panel of cytokines and chemokines in astrocyte conditioned medium. This showed upregulation of a small number of cytokines including CXCL1, M-CSF and CCL2 in medium from TGCM-treated mouse primary astrocytes relative to WTCM treated cells (Fig. 4H-I), and CXCL1 and IP-10 from TGCM-treated iNPC-astro (Fig. 4J-K). Aβ-containing medium significantly increased the secretion of CXCL1 from both mouse and human astrocytes (Fig. 4I, K). Notably, CXCL1 levels are increased in AD brain homogenate relative to samples from age-matched controls (Fig. 4L-M), and CXCL1 has previously been implicated in AD-associated tau changes [56].

**Figure 4:**
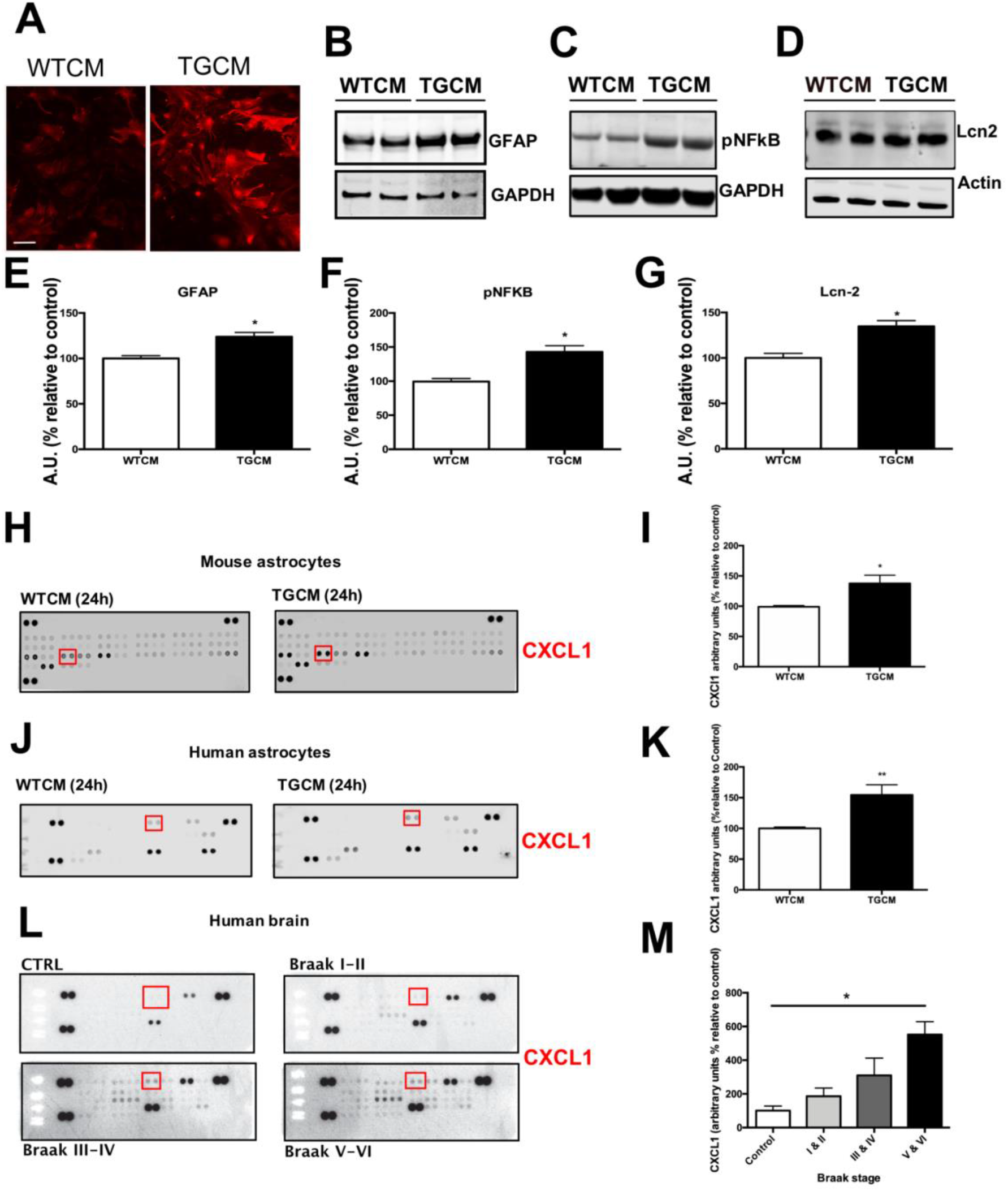
Astrocytic inflammatory phenotypes are induced by Aβ. To explore the effects of physiological Aβ concentrations on astrocyte phenotypes, a number of inflammatory markers were examined. A) Mouse astrocytes exposed to TGCM for 24 hours showed increased GFAP immunoreactivity (red) relative to those treated with WTCM (n= 5). Scale bar = 100μm. Lysates from treated astrocytes were immunoblotted with antibodies against B) GFAP, C) phospho-NFkB (p65) and D) Lcn2. GAPDH was used as a loading control. Quantification of western blot intensities showed increases in E) GFAP, F) pNFkB and G) Lcn2 relative to astrocytes treated with WTCM. Levels of the protein of interest were normalised to GAPDH in each case (n= 5). Antibody-based cytokine arrays were used to provide unbiased analysis of cytokine content in medium from WTCM and TGCM exposed mouse and human astrocytes and in tissue homogenates from postmortem AD and control brain. Representative membranes showing increased CXCL1 in TGCM conditions are shown for H) mouse astrocytes (n= 4) and J) human iNPC-astrocytes (n= 3). L) total protein normalised cytosolic fractions of postmortem control and BA9 prefrontal cortex of AD brain at different Braak stages (n= 4 control, n= 7 stage I-II, n= 10 stage III-IV, n= 6 stage V-VI). Quantification of CXCL1 amounts in these samples revealed significant increases in CXCL1 in TGCM-treated I) mouse and K) human astrocytes, and M) in Braak stage V-VI human AD brain. Data on graphs is mean +/− SEM. *p<0.05.

### CXCL1 is synaptotoxic

To determine if the synaptotoxic effects of culture medium from TGCM-stimulated astrocytes is mediated by CXCL1, we applied recombinant CXCL1 to primary neurons. This significantly reduced spine number by 66.56+/−9.89% (Fig. 5A,B), similar to the reductions identified previously using whole TGCM astro. Measures of neuron health including neurite length and number of nodes and extremities were also affected by recombinant CXCL1 (Fig. 5C-E). CXCL1-mediated synaptotoxicity was prevented when neurons were pre-treated with an antagonist (SB225) of the CXCL1 receptor, CXCR2 (Fig. 5A-E). Notably, direct application of CXCL1 to neurons also replicated the altered localisation of tau observed with TGCM astro, and this was also prevented upon CXCR2 antagonism (Fig. 5F). These results strongly implicate CXCL1 in the synaptotoxic effects of astrocytes in response to physiological concentrations of human Aβ.

**Figure 5:**
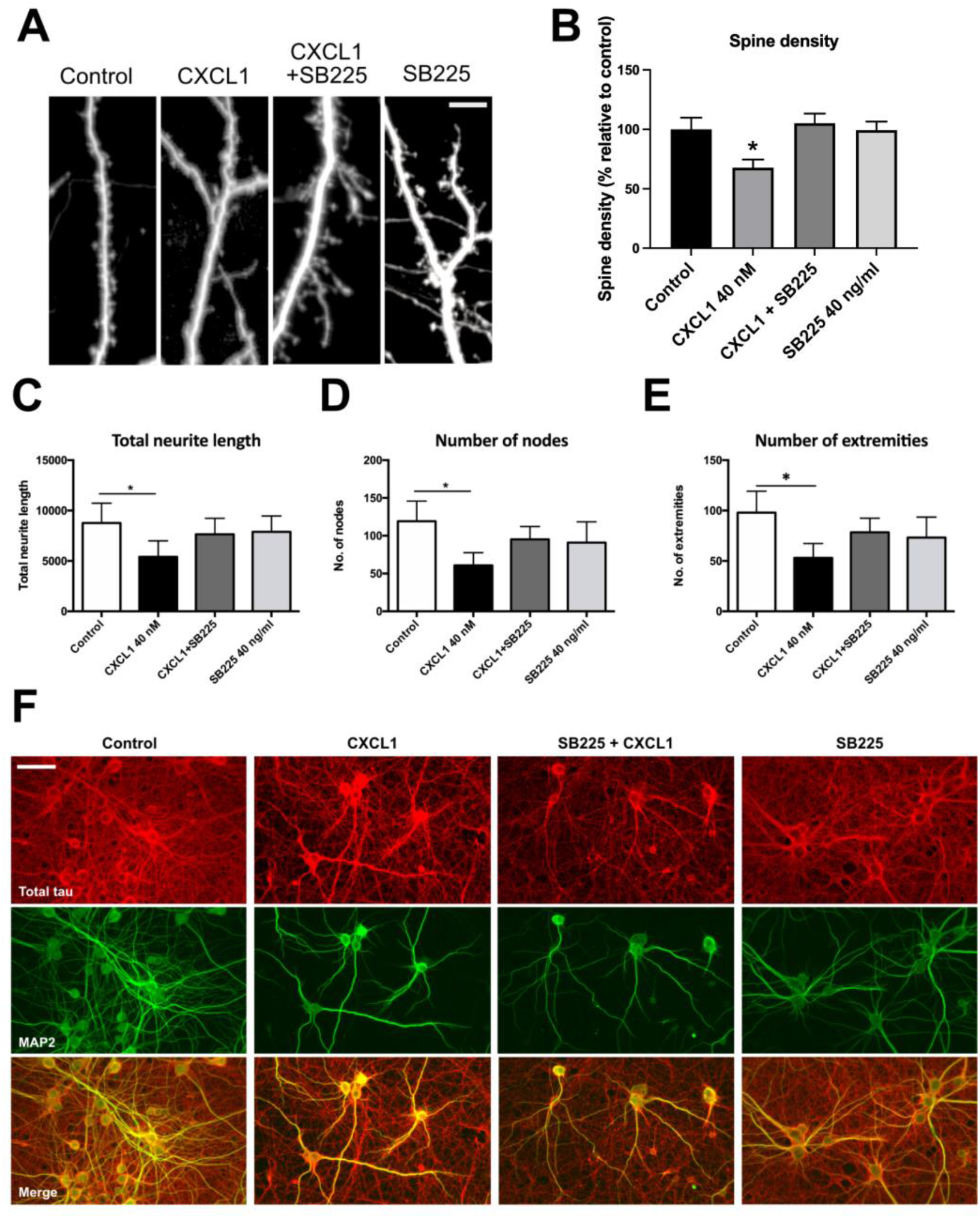
CXCL1 is synaptotoxic and induces tau mislocalisation. Mouse neurons, transfected at 5-7DIV with GFP, were treated with 40 nM recombinant mouse CXCL1 or vehicle for 24 hours to determine if CXCL1 is damaging to neurons. In some cases, neurons were pre-treated for 20 minutes with 40 ng/ml SB225002, an antagonist of the CXCL1 receptor, CXCR2. Neurons were imaged at 14DIV. A) Representative images of dendrites in CXCL1 and SB225002 treated conditions (n= 3) Scale bar = 10μm. B) Quantification of spine density per μm shows reduced spine numbers upon exposure to CXCL1 relative to controls, that was rescued when neurons were pre-treated with the CXCR2 antagonist (n= 6). Measures of neuron health across three wells per experimental repeat showed C) reduced neurite length, D) fewer nodes, and E) fewer extremities on exposure of neurons to CXCL1 that is prevented by pre-treatment with SB225002, demonstrating impairments in neuron health as a result of CXCL1 (n= 3). Data on graphs is mean +/− SEM. *p<0.05. F) Neurons were fixed and immunolabelled with antibodies against MAP2 (green) and tau (red). Exposure to CXCL1 induced increased tau localisation in neurites relative to control conditions that was prevented by antagonism of CXCR2. Scale bar is 50μm.

### CXCL1-CXCR2 interactions mediate the Aβ-induced synaptotoxic responses of astrocytes

Finally, we determined that blocking the CXCR2 receptor is sufficient to prevent synaptotoxicity in response to secretions from Aβ-stimulated astrocytes. Here, astrocytes were stimulated with TGCM or WTCM as before, and the astrocyte conditioned medium was applied to primary neurons that had been pre-incubated with the CXCR2 antagonist SB225. Blocking the CXCL1 receptor prevented the reductions in spine density (Fig. 6A-B) and measures of neuron health (Fig 6C-E) that resulted from Aβ-stimulated astrocyte medium (TGCM astro). These data strongly implicate CXCL1-CXCR2 interactions in the synaptotoxic effects of astrocytes in response to AD-mimicking concentrations of human Aβ.

**Figure 6:**
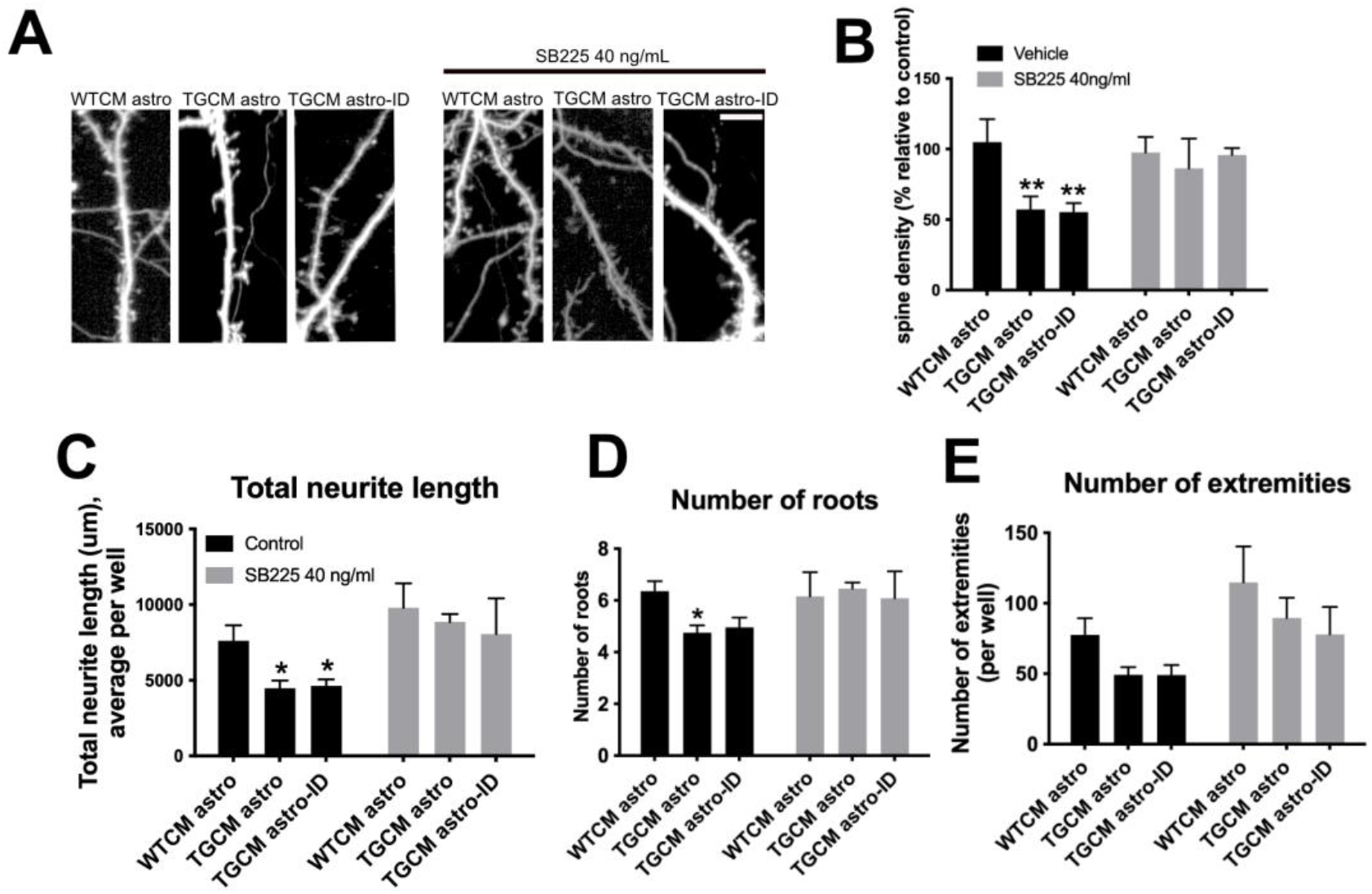
CXCL1-CXCR2 interactions mediate the Aβ-induced synaptotoxic responses of astrocytes. Mouse neurons, transfected at 7DIV with GFP, were exposed to WTCM astro, TGCM astro or TGCM astro-ID for 24 hours in the presence or absence of pre-treatment for 20 minutes with 40 ng/ml SB225002. Neurons were imaged at 14DIV. A) Representative images of dendritic spines (n= 4). Scale bar is 5μm. B) Quantification of spine density per μm showed that reductions in spine numbers upon exposure to TGCM astro and TGCM astro-ID is prevented when neurons are pre-treated with the CXCR2 antagonist. Data is normalised to control conditions (WTCM astro, vehicle). Measures of neuron health across three wells per experimental repeat showed C) reduced neurite length, D) fewer roots, and E) fewer extremities upon exposure of neurons to TGCM astro that is prevented by pre-treatment with SB225002, demonstrating that blocking neuronal CXCR2 prevents the synaptotoxic effects of Aβ-exposed astrocytes (n= 4). Data on graphs is mean +/− SEM. *p<0.05, **p<0.01.

## Discussion

It is now established that non-neuronal cells, and particularly astrocytes and microglia, make major contributions to the onset and progression of Alzheimer’s disease [57–59]. We add novel data to this growing body of evidence to demonstrate that interactions between the astrocytic chemokine CXCL1 and its neuronal receptor CXCR2 promote synaptotoxicity in the presence of Aβ, that is related to tau mislocalisation. As such, our findings further elucidate non-cell autonomous mechanisms underlying synaptotoxic Aβ-tau interactions in AD. There is considerable regional heterogeneity of astrocytes, along with temporal alterations in astrocyte biology with aging and during disease [27, 60] and as such the precise contribution of reactive astrocytes to AD is not clearly established. Our work supports the assertion that rather than becoming reactive secondary to neuronal damage, at least some sub-types of astrocytes respond to AD-mimicking conditions during early “cellular phases” of disease.

Dendritic spines were used here as a measure of synapse health. These structures are post-synaptic sites for the majority of excitatory neurons [61]. Loss of spine density is linked with cognitive decline in humans and is a strong correlate of dementia in AD [1, 62, 63]. Importantly, reductions in spine density in AD are associated with abnormal tau, but not Aβ pathology [62], at least in the prefrontal cortex. However, particularly oligomeric forms of Aβ induce network excitability and synaptoxicity *in vitro* and *in vivo* [64–67]. Together, these findings suggest that while Aβ may drive synaptic dysfunction in AD, tau is the executioner. In keeping with these data, our data show that the loss of dendritic spines upon exposure to TGCM astro is associated with the damaging mis-sorting of tau from the soma.

Tau mislocalisation from the cytoplasmic to synaptic fraction of AD brain is closely correlated with dementia in AD [15] and is a key pathological observation in tauopathy brain [68]. Although some tau is found in dendrites under physiological conditions [69, 70], dendritic tau is increased by Aβ, and can interact with post-synaptic components to further mediate excitotoxicity to Aβ [11, 71].

We add to these findings by showing that in addition to having direct effects on neurons, secretions from Aβ-stimulated astrocytes also induce tau mislocalisation and loss of dendritic spines. “A1” astrocytes were defined by [30] following gene expression analysis from Zamanian et al. [38] as reactive astrocytes with neurotoxic properties. Neurons cultured with “A1” astrocytes, exhibit synaptotoxicity attributed to loss of physiological functions of astrocytes in synaptic maintenance and neuronal homeostasis, alongside secretion of at least one neurotoxic factor [30, 72].

Our data show that astrocytes stimulated by Aβ, which is similar in concentration and species to that found in AD brain [41, 50, 51], increase their release of several cytokines, including the inflammatory chemokine CXCL1. Similar findings have been reported in mouse models of AD and prion disease, particularly following a secondary challenge [36]. We provide evidence that CXCL1 is likely one of the neurotoxic factors secreted by astrocytes since direct application of recombinant CXCL1 recapitulated the loss of dendritic spines that occurs in response to whole conditioned medium from Aβ-stimulated astrocytes (TGCM astro). Importantly, pharmacological blockade of the receptor for CXCL1, CXCR2, prevented both CXCL1- and TGCM astro- induced loss of dendritic spines.

CXCR2 is predominantly expressed in neurons, where it is upregulated proximal to amyloid plaques in AD brain [56]. Importantly, we showed that CXCR2 antagonism was sufficient to prevent CXCL1-induced tau mislocalisation. Indeed, CXCL1 has previously been shown to induce tau phosphorylation via downstream effects on ERK1/2 and PI-3 kinase [56]. CXCL1 also promotes caspase-3 activation [73]. Caspase-3 activity is increased by Aβ, cleaves tau into pro-aggregatory fragments that seed neurofibrillary pathology [74] and is closely linked with synaptic disruption in AD [52]. Direct application of CXCL1 to long-term cultured neurons led to caspase-3 activation, caspase-3 mediated tau cleavage, and tau mislocalisation into bead-like varicosities along neuronal processes [73], as found here. Given the strong links between tau mislocalisation and spine loss, our data strongly support further investigation of the CXCL1-CXCR2 interaction in rodent models of AD with both Aβ and tau abnormalities to determine if this ligand-receptor pair is a novel target for therapeutic intervention.

## Acknowledgements

We thank Dr George Chennell at the Wohl Cellular Imaging Centre at King’s College London for help with microscopy, Professor Dame Pamela Shaw (Sheffield Institute of Translational Neuroscience) for access to fibroblasts from non-ALS controls, and the sadly missed Professor Peter Davies (Feinstein Institute of Medical Research, NY) for the kind gift of tau antibodies. This work was supported by Alzheimer’s Research UK (King’s College London ARUK Network Centre, ARUK-RF2014-2 to BGP-N, ARUK-EG2013-B1 to WN, ARUK-PhD2018-002 to WN and BGP-N, PG2019A-003 to SBW), the Van Geest Charitable Trust (to BGP-N), a grant from Consejo Nacional de Ciencia y Technologia (CONACyT), Mexico (to IV), and grants from the Thierry Latran Foundation and the Academy of Medical Sciences (Ref: SBF002/1142 to LF). Human tissue samples were provided by the London Neurodegenerative Diseases Brain Bank, which receives funding from the MRC and through the Brains for Dementia Research programme (jointly funded by Alzheimer’s Society and Alzheimer’s Research UK).

## Author Contributions

BGP-N and WN designed the study. BGP-N and WN wrote the manuscript. BGP-N performed most experiments and analysed data. LJ, PBL, MMH, LG, FT, MAM, IV-V, MJS, CT, SBW and LF performed additional experiments and/or data analysis and/or contributed to study design. All authors edited and approved the manuscript.

## Data Availability

Summary data are available from the authors on request.

## Supplementary data

**Supplementary Figure 1:**
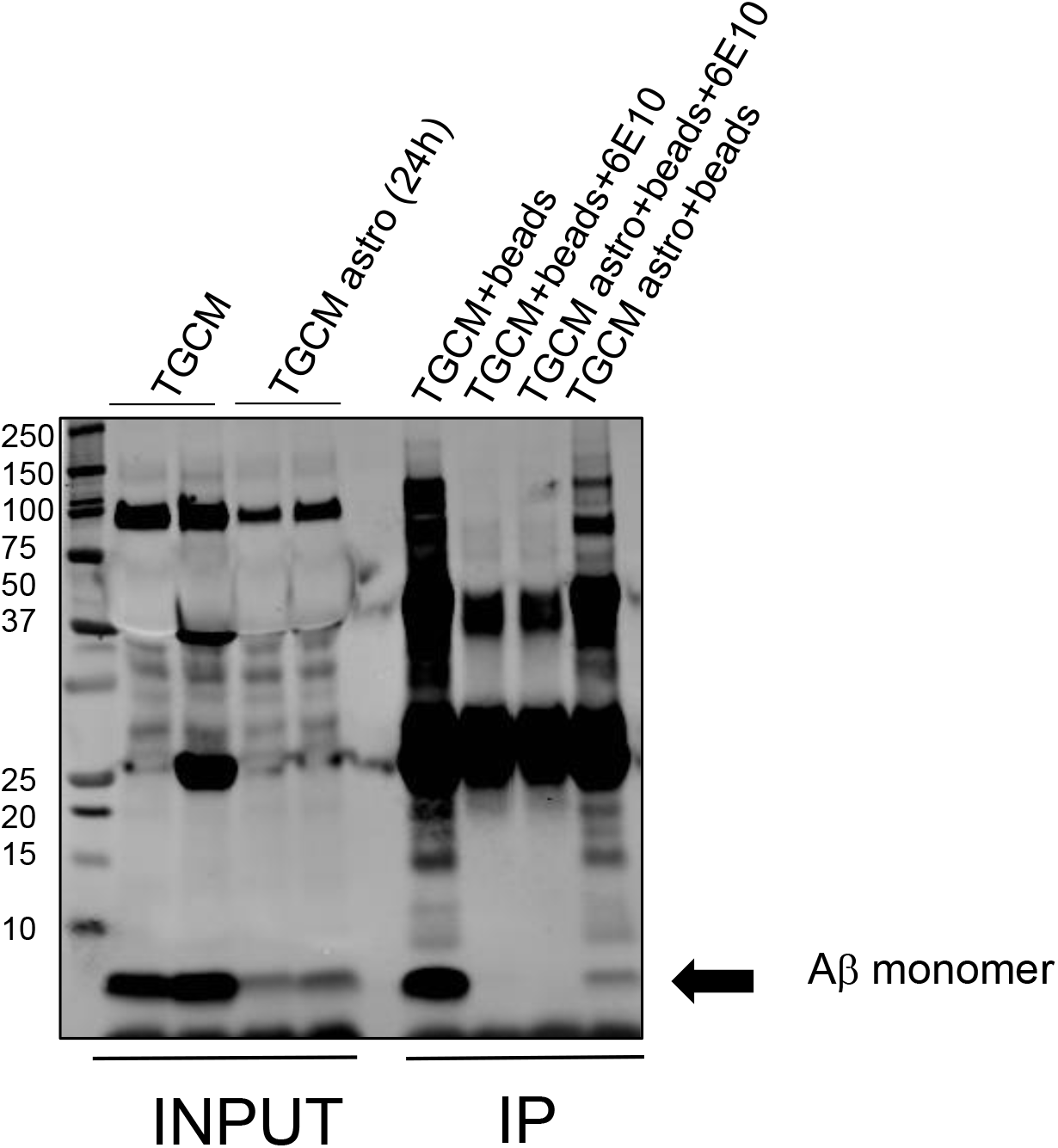
Immunodepletion of Aβ from astrocyte medium with 6E10. Culture medium from astrocytes exposed to TGCM was incubated with protein G Dynabeads (ThermoFisher) bound with 6E10 antibody (COVANCE) for 1 hour at 4°C. Beads were separated from medium using a magnetic stand to remove Aβ-6E10 complexes. Samples of medium before (INPUT) and after immunodepletion were immunoblotted with the 6E10 antibody against Aβ following concentration by immunoprecipitation (IP), showing efficient depletion of Aβ from the culture medium, as indicated by reduced levels of Aβ monomer (black arrow), and other Aβ species.

**Supplementary Figure 2:**
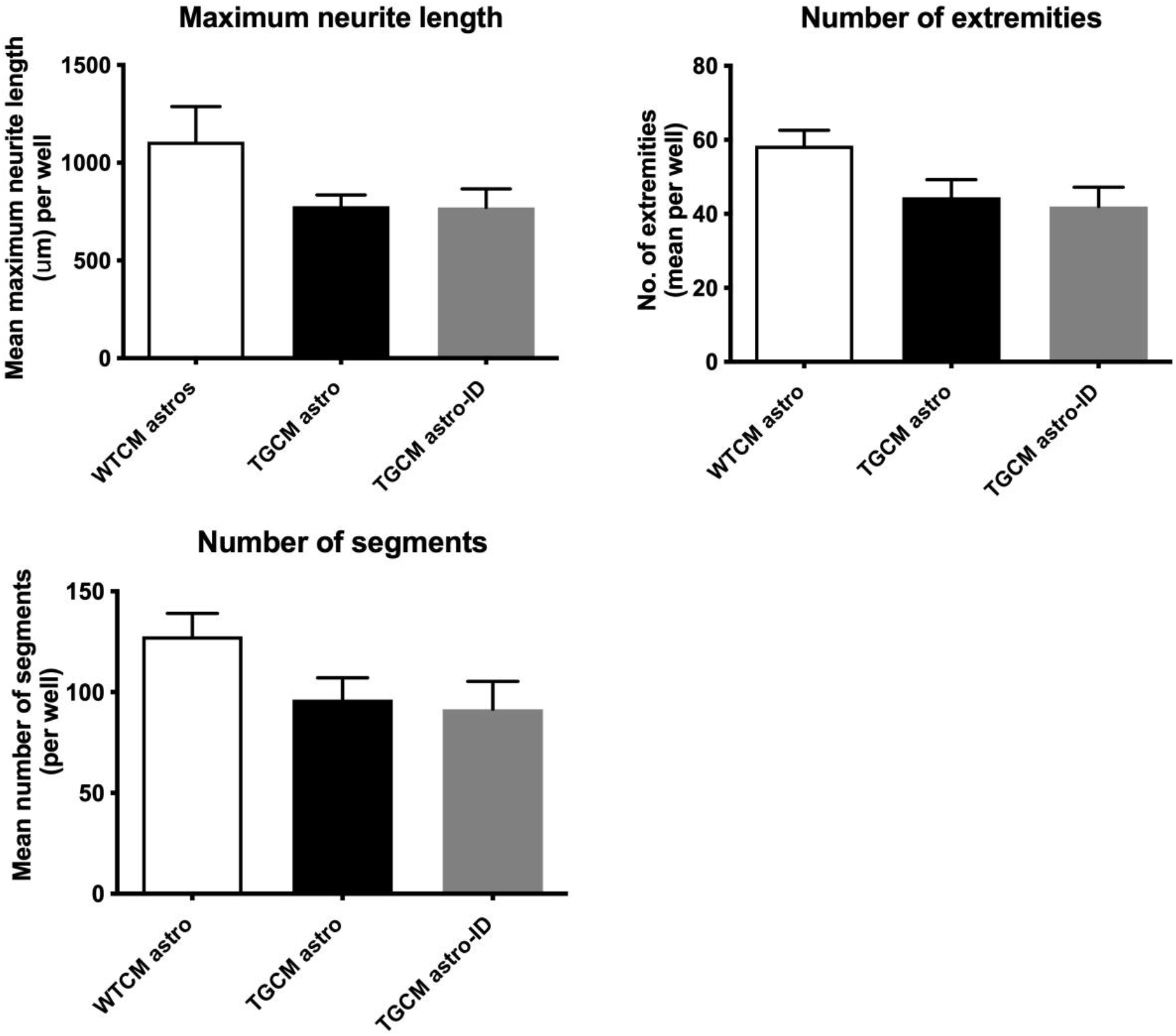
Measures of neuronal health and complexity in mouse neurons challenged with WTCM astro, TGCM astro and TGCM astro-ID. Conditioned medium was collected from primary cortical neurons from wild-type or Tg2576 mice (WTCM; TGCM) and applied to mouse astrocytes. The conditioned medium from stimulated astrocytes was collected, (WTCM astro; TGCM astro) and in some experiments Aβ was immunodepleted from the medium of TGCM treated astrocytes (TGCM astro-ID). These media were added to naïve WT primary neurons that were transfected with GFP at 7DIV. Neuronal complexity as a measure of neuron health was analysed for all cells in three wells per experimental replicate using Harmony software when neurons were 14DIV. This showed non-significant reductions in maximum neurite length, number of extremities and segments following treatment with TGCM astro and TGCM astro-ID (n= 5). Data are mean +/−SEM.

**Supplementary Figure 3:**
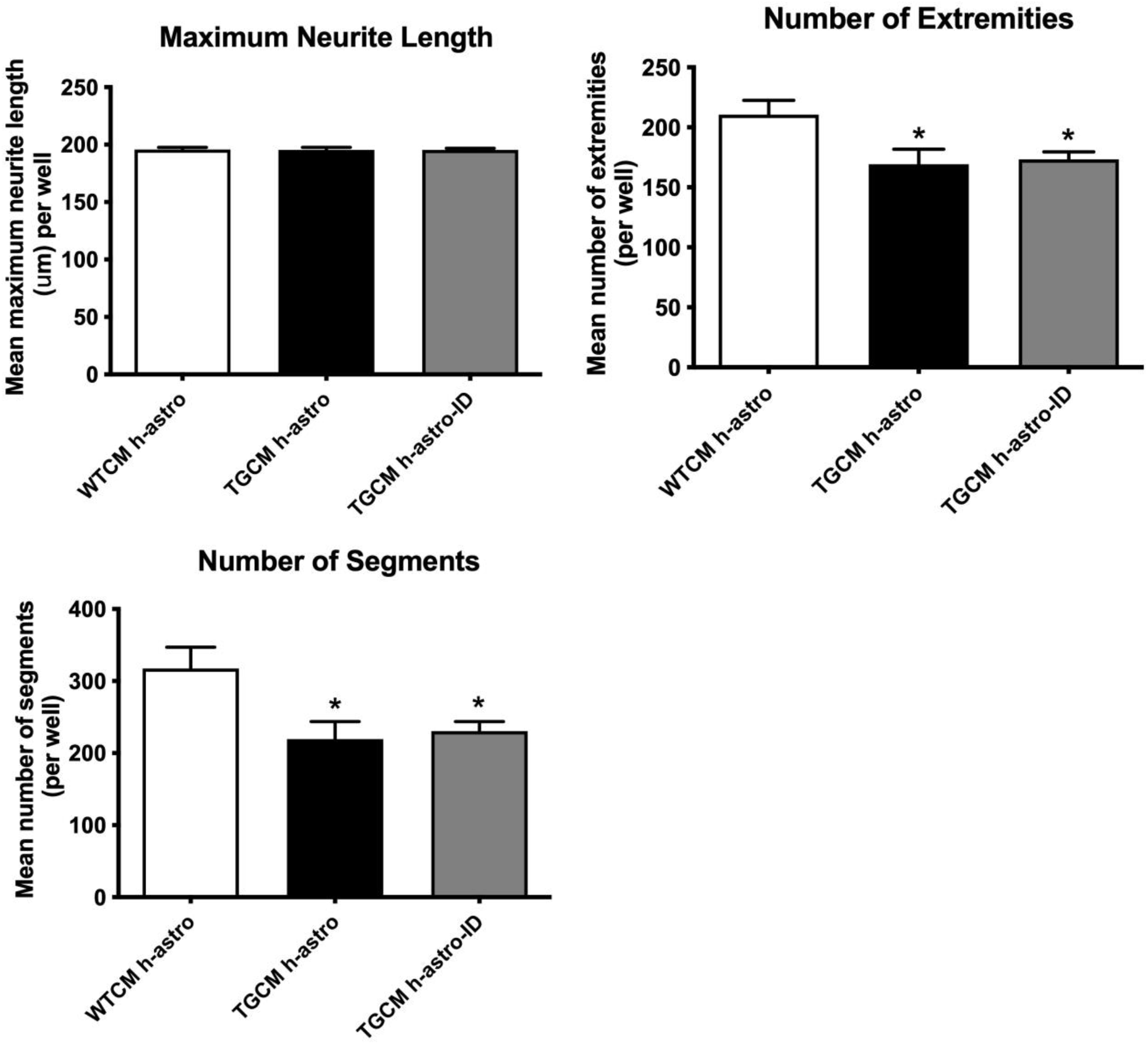
Measures of neuronal health and complexity in LUHMES challenged with WTCM h-astro, TGCM h-astro and TGCM h-astro-ID. Conditioned medium was collected from primary cortical neurons from wild-type or Tg2576 mice (WTCM; TGCM) and applied to human iNPC-astrocytes. The conditioned medium from stimulated astrocytes was collected, (WTCM h-astro; TGCM h-astro) and in some experiments Aβ was immunodepleted from the medium of TGCM treated astrocytes (TGCM h-astro-ID). These media were added to differentiated LUHMES that had been transduced with GFP prior to differentiation. Neuronal complexity as a measure of neuron health was analysed for all cells in three wells per experimental replicate using Harmony software. This showed small reductions in maximum neurite length (not significant), and significant reductions in the number of extremities and segments (p<0.05 for all) following treatment with TGCM h-astro and TGCM h-astro-ID (n= 3). Data are mean +/−SEM.

## References

1. DeKosky, S.T. and S.W. Scheff, Synapse loss in frontal cortex biopsies in Alzheimer’s disease: correlation with cognitive severity. Ann Neurol, 1990. 27(5): p. 457–64.

2. Crimins, J.L., et al., The intersection of amyloid beta and tau in glutamatergic synaptic dysfunction and collapse in Alzheimer’s disease. Ageing Res Rev, 2013. 12(3): p. 757–63.

3. Bertoni-Freddari, C., et al., Synaptic structural dynamics and aging. Gerontology, 1996. 42(3): p. 170–80.

4. Colom-Cadena, M., et al., The clinical promise of biomarkers of synapse damage or loss in Alzheimer’s disease. Alzheimers Res Ther, 2020. 12(1): p. 21.

5. Cleary, J.P., et al., Natural oligomers of the amyloid-beta protein specifically disrupt cognitive function. Nat Neurosci, 2005. 8(1): p. 79–84.

6. Walsh, D.M., et al., Naturally secreted oligomers of amyloid beta protein potently inhibit hippocampal long-term potentiation in vivo. Nature, 2002. 416(6880): p. 535–9.

7. Walsh, D.M., et al., Certain inhibitors of synthetic amyloid beta-peptide (Abeta) fibrillogenesis block oligomerization of natural Abeta and thereby rescue long-term potentiation. J Neurosci, 2005. 25(10): p. 2455–62.

8. Shankar, G.M., et al., Natural oligomers of the Alzheimer amyloid-beta protein induce reversible synapse loss by modulating an NMDA-type glutamate receptor-dependent signaling pathway. J Neurosci, 2007. 27(11): p. 2866–75.

9. Shankar, G.M., et al., Amyloid-beta protein dimers isolated directly from Alzheimer’s brains impair synaptic plasticity and memory. Nat Med, 2008. 14(8): p. 837–42.

10. Walsh, S., et al., Aducanumab for Alzheimer’s disease? BMJ, 2021. 374: p. n1682.

11. Ittner, L.M., et al., Dendritic function of tau mediates amyloid-beta toxicity in Alzheimer’s disease mouse models. Cell, 2010. 142(3): p. 387–97.

12. Hoover, B.R., et al., Tau mislocalization to dendritic spines mediates synaptic dysfunction independently of neurodegeneration. Neuron, 2010. 68(6): p. 1067–81.

13. Zhou, L., et al., Tau association with synaptic vesicles causes presynaptic dysfunction. Nat Commun, 2017. 8: p. 15295.

14. McInnes, J., et al., Synaptogyrin-3 Mediates Presynaptic Dysfunction Induced by Tau. Neuron, 2018. 97(4): p. 823–835 e8.

15. Perez-Nievas, B.G., et al., Dissecting phenotypic traits linked to human resilience to Alzheimer’s pathology. Brain, 2013. 136(Pt 8): p. 2510–26.

16. Augusto-Oliveira, M., et al., Astroglia-specific contributions to the regulation of synapses, cognition and behaviour. Neurosci Biobehav Rev, 2020. 118: p. 331–357.

17. Parpura, V., et al., Glutamate-mediated astrocyte-neuron signalling. Nature, 1994. 369(6483): p. 744–7.

18. Parpura, V. and A. Verkhratsky, The astrocyte excitability brief: from receptors to gliotransmission. Neurochem Int, 2012. 61(4): p. 610–21.

19. Verkhratsky, A., et al., Physiology of Astroglia. Adv Exp Med Biol, 2019. 1175: p. 45–91.

20. Volterra, A. and J. Meldolesi, Astrocytes, from brain glue to communication elements: the revolution continues. Nat Rev Neurosci, 2005. 6(8): p. 626–40.

21. Santello, M., N. Toni, and A. Volterra, Astrocyte function from information processing to cognition and cognitive impairment. Nat Neurosci, 2019. 22(2): p. 154–166.

22. Mederos, S., C. Gonzalez-Arias, and G. Perea, Astrocyte-Neuron Networks: A Multilane Highway of Signaling for Homeostatic Brain Function. Front Synaptic Neurosci, 2018. 10: p. 45.

23. Lushnikova, I., et al., Synaptic potentiation induces increased glial coverage of excitatory synapses in CA1 hippocampus. Hippocampus, 2009. 19(8): p. 753–62.

24. Itagaki, S., et al., Relationship of microglia and astrocytes to amyloid deposits of Alzheimer disease. J Neuroimmunol, 1989. 24(3): p. 173–82.

25. Perez-Nievas, B.G. and A. Serrano-Pozo, Deciphering the Astrocyte Reaction in Alzheimer’s Disease. Front Aging Neurosci, 2018. 10: p. 114.

26. Serrano-Pozo, A., et al., Neuropathological alterations in Alzheimer disease. Cold Spring Harb Perspect Med, 2011. 1(1): p. a006189.

27. Escartin, C., et al., Reactive astrocyte nomenclature, definitions, and future directions. Nat Neurosci, 2021. 24(3): p. 312–325.

28. Bellaver, B., et al., Astrocyte Biomarkers in Alzheimer Disease: A Systematic Review and Meta-analysis. Neurology, 2021.

29. Elahi, F.M., et al., Plasma biomarkers of astrocytic and neuronal dysfunction in early- and late-onset Alzheimer’s disease. Alzheimers Dement, 2020. 16(4): p. 681–695.

30. Liddelow, S.A., et al., Neurotoxic reactive astrocytes are induced by activated microglia. Nature, 2017. 541(7638): p. 481–487.

31. Bhat, R., et al., Astrocyte senescence as a component of Alzheimer’s disease. PLoS One, 2012. 7(9): p. e45069.

32. Garwood, C.J., et al., Astrocytes are important mediators of Abeta-induced neurotoxicity and tau phosphorylation in primary culture. Cell Death Dis, 2011. 2: p. e167.

33. Lian, H., et al., Astrocyte-Microglia Cross Talk through Complement Activation Modulates Amyloid Pathology in Mouse Models of Alzheimer’s Disease. J Neurosci, 2016. 36(2): p. 577–89.

34. Staurenghi, E., et al., Oxysterols present in Alzheimer’s disease brain induce synaptotoxicity by activating astrocytes: A major role for lipocalin-2. Redox Biol, 2021. 39: p. 101837.

35. Cunningham, C., A. Dunne, and A.B. Lopez-Rodriguez, Astrocytes: Heterogeneous and Dynamic Phenotypes in Neurodegeneration and Innate Immunity. Neuroscientist, 2019. 25(5): p. 455–474.

36. Lopez-Rodriguez, A.B., et al., Acute systemic inflammation exacerbates neuroinflammation in Alzheimer’s disease: IL-1beta drives amplified responses in primed astrocytes and neuronal network dysfunction. Alzheimers Dement, 2021.

37. McAlpine, C.S., et al., Astrocytic interleukin-3 programs microglia and limits Alzheimer’s disease. Nature, 2021. 595(7869): p. 701–706.

38. Zamanian, J.L., et al., Genomic analysis of reactive astrogliosis. J Neurosci, 2012. 32(18): p. 6391–410.

39. Zeisel, A., et al., Molecular Architecture of the Mouse Nervous System. Cell, 2018. 174(4): p. 999–1014 e22.

40. Schildge, S., et al., Isolation and culture of mouse cortical astrocytes. J Vis Exp, 2013(71).

41. Hudry, E., et al., Inhibition of the NFAT pathway alleviates amyloid beta neurotoxicity in a mouse model of Alzheimer’s disease. J Neurosci, 2012. 32(9): p. 3176–92.

42. Arbel-Ornath, M., et al., Soluble oligomeric amyloid-beta induces calcium dyshomeostasis that precedes synapse loss in the living mouse brain. Mol Neurodegener, 2017. 12(1): p. 27.

43. Meyer, K., et al., Direct conversion of patient fibroblasts demonstrates non-cell autonomous toxicity of astrocytes to motor neurons in familial and sporadic ALS. Proc Natl Acad Sci U S A, 2014. 111(2): p. 829–32.

44. Allen, S.P., et al., Astrocyte adenosine deaminase loss increases motor neuron toxicity in amyotrophic lateral sclerosis. Brain, 2019. 142(3): p. 586–605.

45. Varcianna, A., et al., Micro-RNAs secreted through astrocyte-derived extracellular vesicles cause neuronal network degeneration in C9orf72 ALS. EBioMedicine, 2019. 40: p. 626–635.

46. Ratcliffe, L.E., et al., Loss of IGF1R in Human Astrocytes Alters Complex I Activity and Support for Neurons. Neuroscience, 2018. 390: p. 46–59.

47. Croft, C.L. and W. Noble, Preparation of organotypic brain slice cultures for the study of Alzheimer’s disease. F1000Res, 2018. 7: p. 592.

48. Glennon, E.B., et al., Bridging Integrator-1 protein loss in Alzheimer’s disease promotes synaptic tau accumulation and disrupts tau release. Brain Commun, 2020. 2(1).

49. Srivastava, D.P., K.M. Woolfrey, and P. Penzes, Analysis of dendritic spine morphology in cultured CNS neurons. J Vis Exp, 2011(53): p. e2794.

50. Wu, H.Y., et al., Distinct dendritic spine and nuclear phases of calcineurin activation after exposure to amyloid-beta revealed by a novel fluorescence resonance energy transfer assay. J Neurosci, 2012. 32(15): p. 5298–309.

51. DaRocha-Souto, B., et al., Activation of glycogen synthase kinase-3 beta mediates beta-amyloid induced neuritic damage in Alzheimer’s disease. Neurobiol Dis, 2012. 45(1): p. 425–37.

52. D’Amelio, M., M. Sheng, and F. Cecconi, Caspase-3 in the central nervous system: beyond apoptosis. Trends Neurosci, 2012. 35(11): p. 700–9.

53. Li, J., et al., Conservation and divergence of vulnerability and responses to stressors between human and mouse astrocytes. Nat Commun, 2021. 12(1): p. 3958.

54. Gatto, N., et al., Directly converted astrocytes retain the ageing features of the donor fibroblasts and elucidate the astrocytic contribution to human CNS health and disease. Aging Cell, 2021. 20(1): p. e13281.

55. Scholz, D., et al., Rapid, complete and large-scale generation of post-mitotic neurons from the human LUHMES cell line. J Neurochem, 2011. 119(5): p. 957–71.

56. Xia, M. and B.T. Hyman, GROalpha/KC, a chemokine receptor CXCR2 ligand, can be a potent trigger for neuronal ERK1/2 and PI-3 kinase pathways and for tau hyperphosphorylation-a role in Alzheimer’s disease? J Neuroimmunol, 2002. 122(1-2): p. 55–64.

57. Henstridge, C.M., B.T. Hyman, and T.L. Spires-Jones, Beyond the neuron-cellular interactions early in Alzheimer disease pathogenesis. Nat Rev Neurosci, 2019. 20(2): p. 94–108.

58. Scheltens, P., et al., Alzheimer’s disease. Lancet, 2021. 397(10284): p. 1577–1590.

59. Paumier, A., et al., Astrocyte-neuron interplay is critical for Alzheimer’s disease pathogenesis and is rescued by TRPA1 channel blockade. Brain, 2021.

60. Boisvert, M.M., et al., The Aging Astrocyte Transcriptome from Multiple Regions of the Mouse Brain. Cell Rep, 2018. 22(1): p. 269–285.

61. Walker, C.K. and J.H. Herskowitz, Dendritic Spines: Mediators of Cognitive Resilience in Aging and Alzheimer’s Disease. Neuroscientist, 2020: p. 1073858420945964.

62. Boros, B.D., et al., Dendritic spines provide cognitive resilience against Alzheimer’s disease. Ann Neurol, 2017. 82(4): p. 602–614.

63. Chen, M.K., et al., Assessing Synaptic Density in Alzheimer Disease With Synaptic Vesicle Glycoprotein 2A Positron Emission Tomographic Imaging. JAMA Neurol, 2018. 75(10): p. 1215–1224.

64. Blurton-Jones, M. and F.M. Laferla, Pathways by which Abeta facilitates tau pathology. Curr Alzheimer Res, 2006. 3(5): p. 437–48.

65. Ittner, L.M. and J. Gotz, Amyloid-beta and tau--a toxic pas de deux in Alzheimer’s disease. Nat Rev Neurosci, 2011. 12(2): p. 65–72.

66. Mucke, L. and D.J. Selkoe, Neurotoxicity of amyloid beta-protein: synaptic and network dysfunction. Cold Spring Harb Perspect Med, 2012. 2(7): p. a006338.

67. Siskova, Z., et al., Dendritic structural degeneration is functionally linked to cellular hyperexcitability in a mouse model of Alzheimer’s disease. Neuron, 2014. 84(5): p. 1023–33.

68. Braak, H., et al., Stages of the pathologic process in Alzheimer disease: age categories from 1 to 100 years. J Neuropathol Exp Neurol, 2011. 70(11): p. 960–9.

69. Ittner, A. and L.M. Ittner, Dendritic Tau in Alzheimer’s Disease. Neuron, 2018. 99(1): p. 13–27.

70. Xia, D., J.M. Gutmann, and J. Gotz, Mobility and subcellular localization of endogenous, gene-edited Tau differs from that of over-expressed human wild-type and P301L mutant Tau. Sci Rep, 2016. 6: p. 29074.

71. Li, C. and J. Gotz, Somatodendritic accumulation of Tau in Alzheimer’s disease is promoted by Fyn-mediated local protein translation. EMBO J, 2017. 36(21): p. 3120–3138.

72. Guttenplan, K.A., et al., Knockout of reactive astrocyte activating factors slows disease progression in an ALS mouse model. Nat Commun, 2020. 11(1): p. 3753.

73. Zhang, X.F., et al., CXCL1 Triggers Caspase-3 Dependent Tau Cleavage in Long-Term Neuronal Cultures and in the Hippocampus of Aged Mice: Implications in Alzheimer’s Disease. J Alzheimers Dis, 2015. 48(1): p. 89–104.

74. Cotman, C.W., et al., The role of caspase cleavage of tau in Alzheimer disease neuropathology. J Neuropathol Exp Neurol, 2005. 64(2): p. 104–12.

